# PDBspheres - a method for finding 3D similarities in local regions in proteins

**DOI:** 10.1101/2022.01.04.474934

**Authors:** Adam T. Zemla, Jonathan E. Allen, Dan Kirshner, Felice C. Lightstone

## Abstract

We present a structure-based method for finding and evaluating structural similarities in protein regions relevant to ligand binding. PDBspheres comprises an exhaustive library of protein structure regions (“spheres”) adjacent to complexed ligands derived from the Protein Data Bank (PDB), along with methods to find and evaluate structural matches between a protein of interest and spheres in the library. PDBspheres uses the LGA structure alignment algorithm as the main engine for detecting structure similarities between the protein of interest and template spheres from the library, which currently contains more than 2 million spheres. To assess confidence in structural matches an all-atom-based similarity metric takes sidechain placement into account. Here, we describe the PDBspheres method, demonstrate its ability to detect and characterize binding sites in protein structures, show how PDBspheres - a strictly structure-based method - performs on a curated dataset of 2,528 ligand-bound and ligand-free crystal structures, and use PDBspheres to cluster pockets and assess structural similarities among protein binding sites of 4,876 structures in the “refined set” of the PDBbind 2019 dataset.

## INTRODUCTION

Interactions between proteins and small molecule ligands are a cornerstone of biochemical function. Modern drug discovery often relies on structural information about the target of interest (typically a protein). However, when a new structure is obtained, it may be the case that little is known with regard to potential binding sites on that structure. Numerous studies have focused on understanding protein-ligand interactions (e.g., 1,2) to further the development of binding site prediction methods (e.g., 3-6) as well as benchmark datasets (e.g., 7-11) used to compare and evaluate their performance. In general, the binding site prediction methods can be categorized in three broad sets: 1) template-based methods that use known protein information, 2) physics-based methods that rely on geometry (for example, cavity detection) and/or physicochemical properties (for example, surface energy interactions with probe molecules), and 3) machine learning (ML) - rapidly developing in recent years methods capable of efficiently processing information collected in their training data sets. For ML methods data can come from both experiments and in silico data processing, as described for example in (12-15). Our method, PDBspheres is template-based; however, it relies solely on local structure conformation similarities with templates extracted from structures from PDB – i.e., PDBspheres does not use any prior information from libraries of sequences, motifs or residues forming binding sites that are collected separately to enhance the method performance. In general, template-based methods can be categorized as 1) those that rely on only protein sequence, e.g., Firestar (16), 2) those that rely on only protein structure, e.g., SP-ALIGN (17), or 3) those that rely both on sequence and structure, e.g., Concavity (18). Some methods, e.g., SURFNET (19) are classified as structure-based methods, but since they use geometric characteristics, they are grouped within geometric-based methods (20).

One of the strengths of the developed PDBspheres method is that in the binding site searches we use only those template models of cavities (PDBspheres library) that are already observed as protein-ligand binding sites in other proteins (experimentally solved protein-ligand complexes deposited in PDB). This approach ensures that any structural shape which might resemble a cavity in evaluated protein models – e.g., holes within interchain interfaces, missing fragments in coordinates, or other structural imperfections in geometry – will not be reported by PDBspheres searches as cavity candidates, simply because they are not evidenced as *ligand-binding* cavities in reality, i.e., in confirmed-by-experiment PDB structures. To resolve the problem with this kind of false-positive pocket assignments some template-based methods may need additional information about proteins collected from calculated sequence alignments, patterns in residues forming binding sites, etc. A natural strength of structure-template-based methods such as PDBspheres lies in their ability to detect similar binding sites and help determine the biochemical function even if the sequence identity is very low, i.e., when similarity between pockets cannot be detected with confidence based on the sequence. A limitation, however, of the PDBspheres method is that its library depends on structural information derived from the PDB database, so identification of a new pocket or function relies on data availability in PDB, i.e., pockets having shapes not represented in PDB will not be found.

In the PDBspheres method, binding sites are identified exclusively based on the structure similarity between regions from the query protein and pocket spheres from the PDBspheres library. Ligand placement within a predicted pocket is calculated based on a protein-sphere superposition, i.e., an agreement in structure conformation between atoms from the query protein and protein atom coordinates from the template sphere. The main premise of structure template-based binding site prediction methods is that the number of pockets is limited (2), therefore each one of them may serve as a binding site for a large diversity of ligands (17). However, possible conformations of ligands bound in the pocket are also limited. Indeed, when we predict a ligand-protein conformation we focus on what parts of the ligand (e.g., its core region) need to be in a correct conformation, i.e., a conformation that can be confirmed by experiment and from which the binding affinities can be estimated. Usually such “correct” conformations are shared between ligands that bind a given pocket (at least shared by their “core” regions.) The evaluation of the correctness of the ligand placement within a pocket is not an easy task. For example, when for a given protein two pocket predictions with different ligands inserted need to be evaluated a simple calculation and comparison of the ligand centroids (the center of mass of the bound ligands) can be misleading. This is because the two predictions may differ in assigned placements of the ligands within a pocket (especially when the pocket size is large allowing the ligands to fit in different areas within the cavity), or the ligand sizes and their shapes are different (e.g., some parts of the ligands can be exposed outside the pocket with different orientations (21); see Figure 1). These difficulties in the assessment of the accuracy of binding site predictions and ligand placements within predicted pockets could be overcome when we focus on the evaluation of residues interacting with the bound ligands. In Figure 1 we present an example of the b1 domain from the Human Neuropilin-1 (PDB 2qqi) where two different poses of the same ligand EG01377 (PDB id: DUE) are possible (21). Superposition of two experimentally solved structures of Neuropilin1-b1 domain in complex with EG01377 shows almost identical protein structure conformations while poses of bound ligands differ significantly outside the “core”. The distance between centroids of the ligand’s two orientations placed in the pocket is more than 4.6 Å when the centroid is calculated on all ligand’s atoms, and only 0.7 Å when it is calculated on the buried parts of the ligands - “core”.

**Figure 1.**
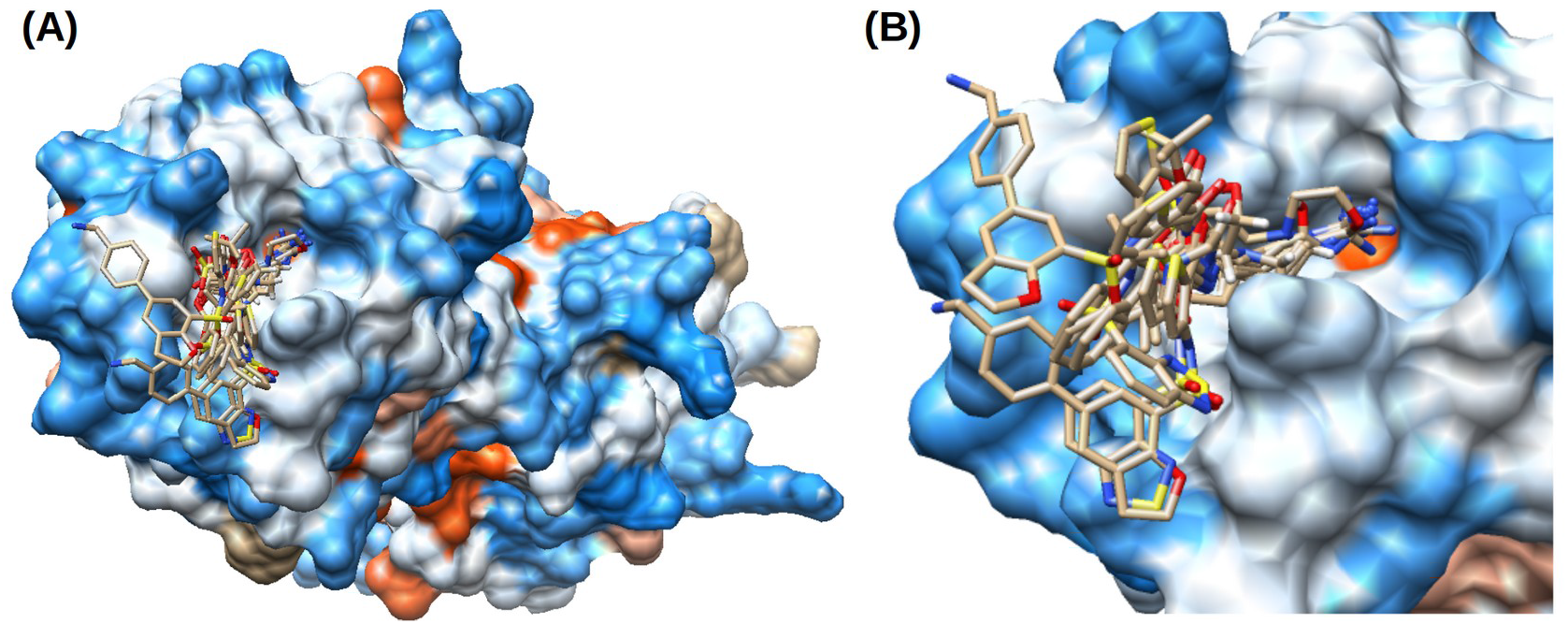
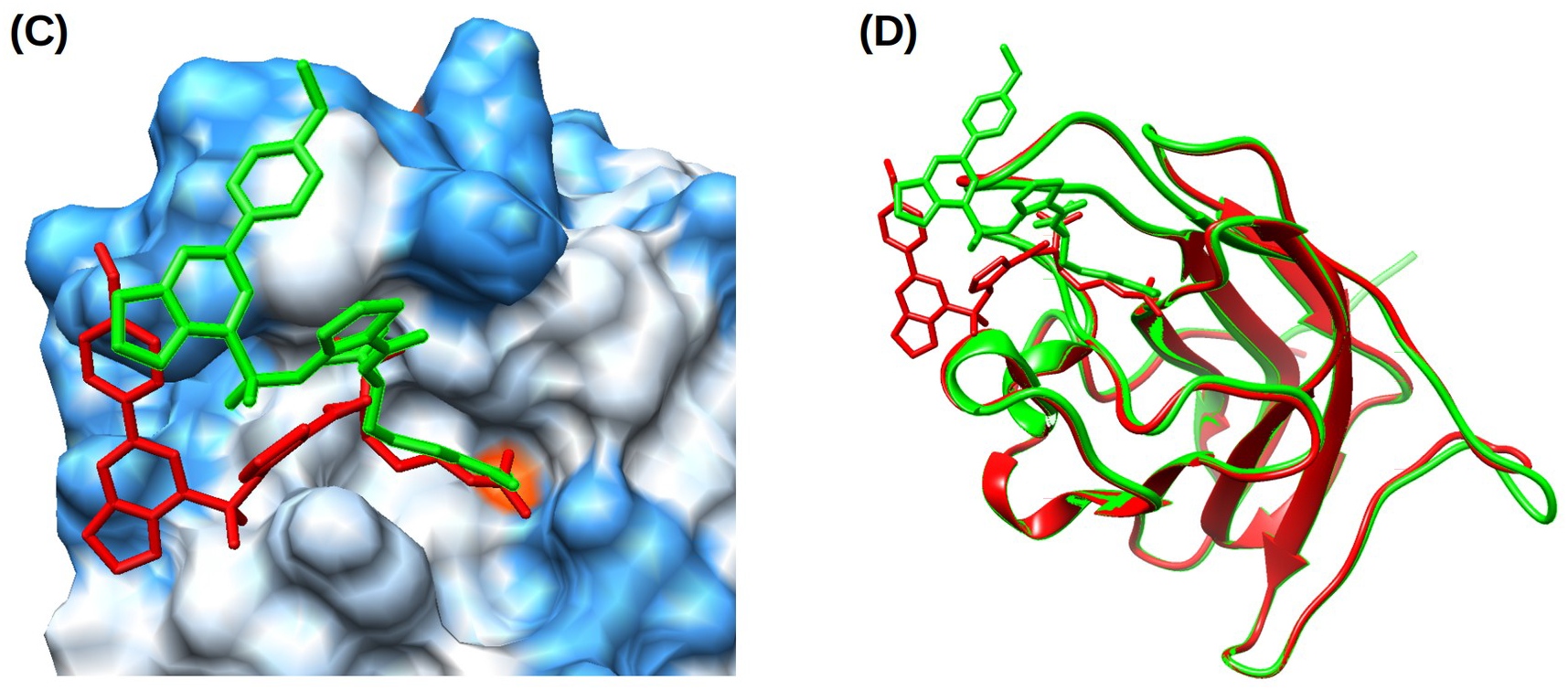
(A) Example of the b1 domain from the Human Neuropilin-1 (PDB 2qqi) where a single pocket can serve as a binding site for many different ligands. (B) Zoom-in to the pocket that can accommodate ligands of different sizes. A list of ligands includes: 6JY.20 (Arg-7), 6K8.24 (Arg-6), 8DR.32 (EG00229), AAG.15 (M45), AR5.19 (Arg-5), BCN.23 (Bicine), DUE.40 (EG01377), HRG.13 (Arg-1), R40.22 (Arg-4), where ligand information “PDBid.size” is provided with a size representing a number of heavy atoms. (C) Two different poses of the ligand DUE.40 (EG01377) are reported in Powell et al. 2018 (21) and identified by PDBspheres based on two templates: 6fmc and 6fmf. (D) Superposition of two experimentally solved structures of Neuropilin1-b1 domain in complex with EG01377 (in green: PDB 6fmc at resolution 0.9 Å, and in red: PDB 6fmf at resolution 2.8 Å) shows almost identical protein structure conformations while poses of bound ligands differ significantly outside the core. The distance between centroids of the ligand’s two orientations when placed in the pocket is more than 4.6 Å.

Knowing the correct “core” conformation of the ligand within the pocket (i.e., the pose of the conserved part) can significantly improve “*in silico*” drug discovery; it can be used in compound screening/docking efforts as a pre-filter for selection of most promising compounds for a more detailed and expensive computational evaluation. Currently, there are millions of compounds screened using docking or molecular mechanics-generalized Born surface area (MMGBSA) approaches. These methods are widely used to estimate ligand-binding affinities in identified binding sites of targeted proteins to find good candidates for a further (experimental) analysis of potential inhibitors.

Assessment of the performance of binding sites prediction methods is not an easy task because they may perform differently on different benchmarks. In Critical Assessment of Protein Structure Prediction (CASP) experiments (rounds 8, 9, and 10) (22-24) an attempt was made to compare different ligand binding residue prediction methods, but it was found that the assessment might be biased (not enough targets, targets not diverse enough to represent different families, folds, ligand type, and ligand binding sites) and this category was subsequently dropped from CASP. A comprehensive benchmark dataset was recently designed by Clark et al (25,26). It covers 304 unique protein families with 2,528 structures in 1456 “holo” and 1082 “apo” conformations. A detailed comparison analysis using this benchmark dataset was performed on seven binding site prediction methods (25). The authors decided to exclude template-based methods from their evaluation to avoid unfair comparison with non-template-based methods. As stated in (25) template-based methods use “libraries of sequence-based template information” which “would inevitably lead to the use of holo structure knowledge to solve the binding site locations in apo structures.” So, it was expected that template-based methods would easily outperform non-template-based methods on the benchmark.

The accuracy of pocket prediction methods in Clark et al. (25) analysis and during CASP was measured mainly by the Mathew’s Correlation Coefficient (MCC). This metric assigns high scores not just when at least one residue in a given binding site is identified correctly, but rather rewards methods with more correct and less false predictions of contact residues implying which of the algorithm pocket predictions are close to the “correct” location on the binding surface of the protein. In this paper, we use the same metric to evaluate the performance of our method with varying datasets of structural templates used by PDBsheres to make predictions. Our reported results were merged with results from the assessment of non-template-based methods performed by Clark et al. (25) in Table 1 and also a discussion of the performance of template-based methods from CASP (22-24) is included to illustrate how PDBspheres places within these methods and what the limits of structure-based pocket prediction methods may be.

**Table 1.**
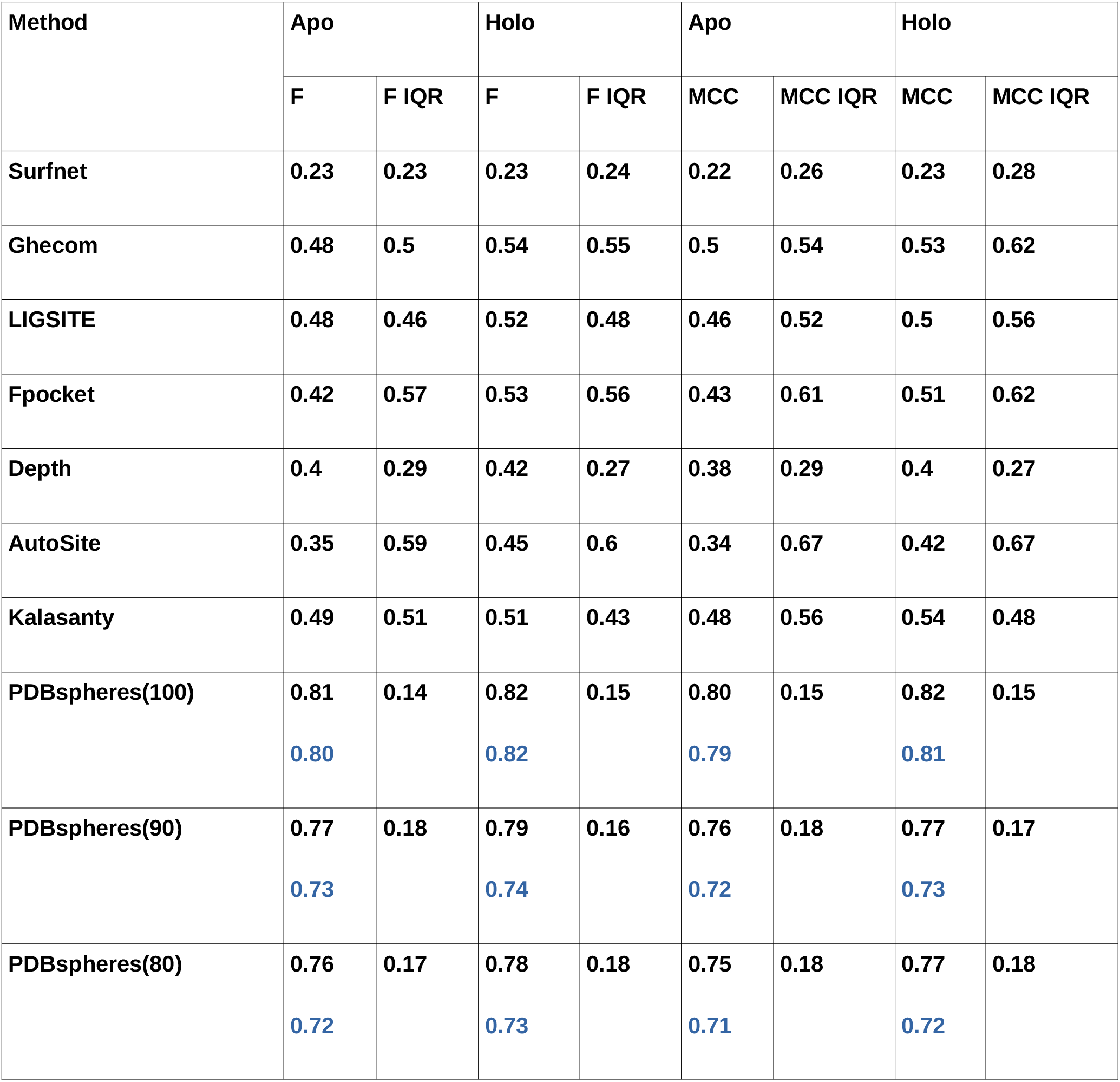
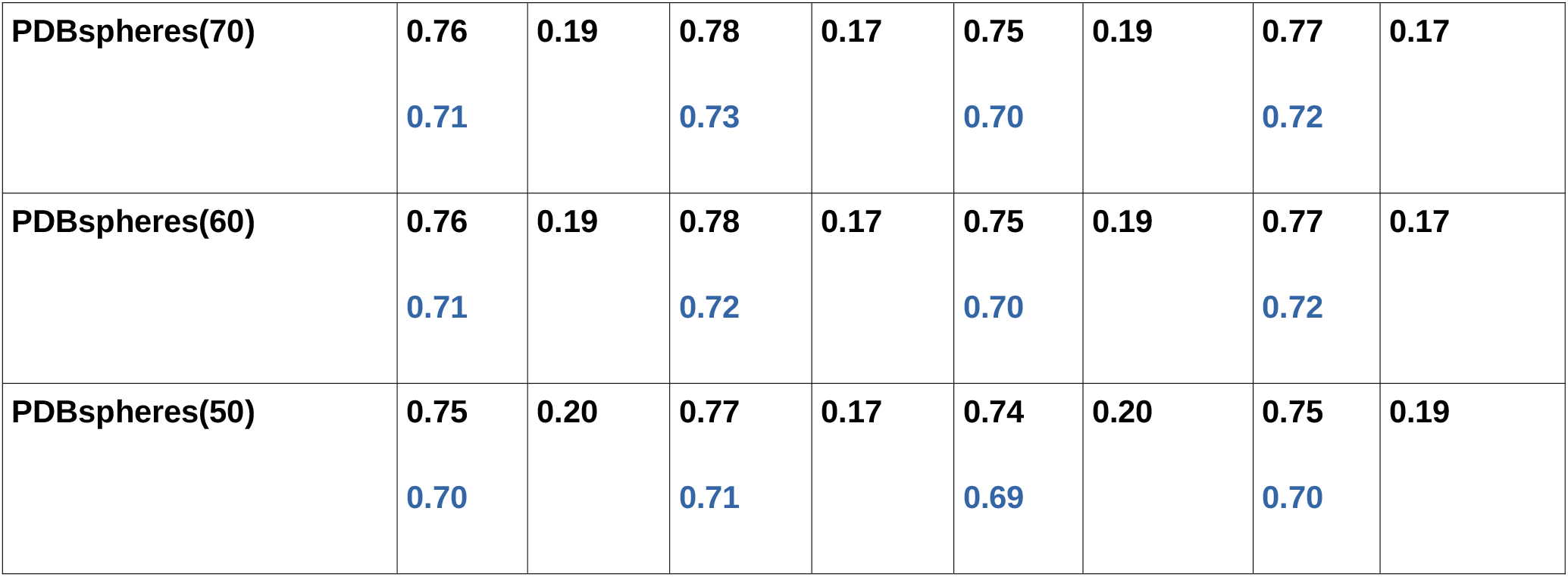
Median of family median F scores and MCCs for apo and holo datasets for all seven “non-template-based” LBS-prediction methods described in Clark et al. (25), and PDBspheres (for which mean values are also given, the second line in the rows for PDBspheres colored in blue). IQR (interquartile range) describes the difference between maximum and minimum scores within the middle 50% of values when ordered from lowest to highest. IQR indicates how close the middle 50% of family F and MCC values are to their respective medians. F and MCC scores are described in Section: Benchmark dataset and evaluation metrics.

| Method | Apo |  | Holo |  | Apo |  | Holo |  |
| --- | --- | --- | --- | --- | --- | --- | --- | --- |
|  | F | F IQR | F | F IQR | MCC | MCC IQR | MCC | MCC IQR |
| Surfnet | 0.23 | 0.23 | 0.23 | 0.24 | 0.22 | 0.26 | 0.23 | 0.28 |
| Ghecom | 0.48 | 0.5 | 0.54 | 0.55 | 0.5 | 0.54 | 0.53 | 0.62 |
| LIGSITE | 0.48 | 0.46 | 0.52 | 0.48 | 0.46 | 0.52 | 0.5 | 0.56 |
| Fpocket | 0.42 | 0.57 | 0.53 | 0.56 | 0.43 | 0.61 | 0.51 | 0.62 |
| Depth | 0.4 | 0.29 | 0.42 | 0.27 | 0.38 | 0.29 | 0.4 | 0.27 |
| AutoSite | 0.35 | 0.59 | 0.45 | 0.6 | 0.34 | 0.67 | 0.42 | 0.67 |
| Kalasanty | 0.49 | 0.51 | 0.51 | 0.43 | 0.48 | 0.56 | 0.54 | 0.48 |
| PDBspheres(100) | 0.81<br>0.80 | 0.14 | 0.82<br>0.82 | 0.15 | 0.80<br>0.79 | 0.15 | 0.82<br>0.81 | 0.15 |
| PDBspheres(90) | 0.77<br>0.73 | 0.18 | 0.79<br>0.74 | 0.16 | 0.76<br>0.72 | 0.18 | 0.77<br>0.73 | 0.17 |
| PDBspheres(80) | 0.76<br>0.72 | 0.17 | 0.78<br>0.73 | 0.18 | 0.75<br>0.71 | 0.18 | 0.77<br>0.72 | 0.18 |
| **PDBspheres(70)** | **0.76**<br>**0.71** | **0.19** | **0.78**<br>**0.73** | **0.17** | **0.75**<br>**0.70** | **0.19** | **0.77**<br>**0.72** | **0.17** |
| **PDBspheres(60)** | **0.76**<br>**0.71** | **0.19** | **0.78**<br>**0.72** | **0.17** | **0.75**<br>**0.70** | **0.19** | **0.77**<br>**0.72** | **0.17** |
| **PDBspheres(50)** | **0.75**<br>**0.70** | **0.20** | **0.77**<br>**0.71** | **0.17** | **0.74**<br>**0.69** | **0.20** | **0.75**<br>**0.70** | **0.19** |

## METHODS

The PDBspheres system is designed to help assess the similarity between proteins based on their structure similarity in selected local regions (e.g., binding sites, protein interfaces, or any other local regions that might be structurally characteristic of a particular group of proteins). The PDBspheres system has three main components: 1) the PDBspheres library of binding site templates, 2) a structure similarity search algorithm to detect similarity between evaluated local structural regions (e.g., binding sites), and 3) numerical metrics to assess confidence in detected similar regions.

Currently, the PDBspheres library (ver. 2021/10/13) contains 2,002,354 compound binding site models (template spheres) and 67,445 short-peptide binding site models derived from protein structures deposited in the PDB database (27).

The Local-Global Alignment (LGA) program (28) is used to perform all structure similarity searches. While comparing two structures, the LGA algorithm generates many different local superpositions to detect regions where proteins are similar, and calculated similarity scores are not affected by some perturbations in small parts of compared proteins. In essence, LGA assesses structure similarity based on detected conserved parts between two proteins or protein fragments while removing small deviating regions from calculated scores. This capability of LGA allows accurate detection of similar pocket shapes. The main metrics to assess confidence in detected pockets are a scoring function LGA_S (a combination of Global Distance Test (GDT) and Longest Continuous Segments (LCS) measures (28)), which evaluates structure similarities on Calpha and/or Cbeta levels, and GDC (Global Distance Calculations (29)), which allows evaluation on an all-atom level, i.e., extending structure similarity evaluation to the conformation of side-chain atoms.

When applied to predict protein binding sites in a given protein structure, PDBspheres detects ligand-protein binding regions using the pocket/sphere templates from the PDBspheres library constructed from all available structures deposited in the PDB database. After a binding pocket is detected the ligand(s) from matching template(s) is/are inserted into the identified pocket in a query protein to illustrate an approximate location of the ligand. The location is approximate because it is based on the alignment of the query protein with the template protein spheres, and no docking or any energy minimization or structure relaxation are undertaken. Thus, it should be noted that PDBspheres is not a docking system to predict *de novo* ligand poses within a binding site. However, further relaxation of the predicted protein-ligand complex can be considered as the next step to improve a ligand placement within a pocket (which is a subject of continuing PDBspheres system development.)

In the process of detecting pocket candidates within protein structures, similarity searches can be performed using all templates in the PDBspheres library (i.e., exhaustive search, testing all 2.0 M pockets from the library), or if time is prohibitive, searches can be performed on a preselected subset of the sphere templates. For example, the preselection process can be based on specific targeted ligands (e.g., their names or sizes), or can be limited to those template spheres from the PDBspheres library that come from homologous proteins. The computational time of the structure processing can vary depending on the size of the protein or the number of template spheres from the PDBspheres library preselected by the system for the pocket detection and similarity evaluation. An exhaustive search (against the entire PDBspheres library) can take more than one day on a single processor machine, however with above-mentioned preselection procedures the calculations can be completed in less than one hour for medium size proteins. For example, the processing of three protein structures discussed in the manuscript and illustrated on Figure 1 (Neuropilin-1, 158aa), Figure 2 (PL2pro, 318aa), and Figure 6 (Tryptase, 248aa)) took 7 min, 52 min, and 50 min respectively on a single processor with 16 cores machine.

**Figure 2.**
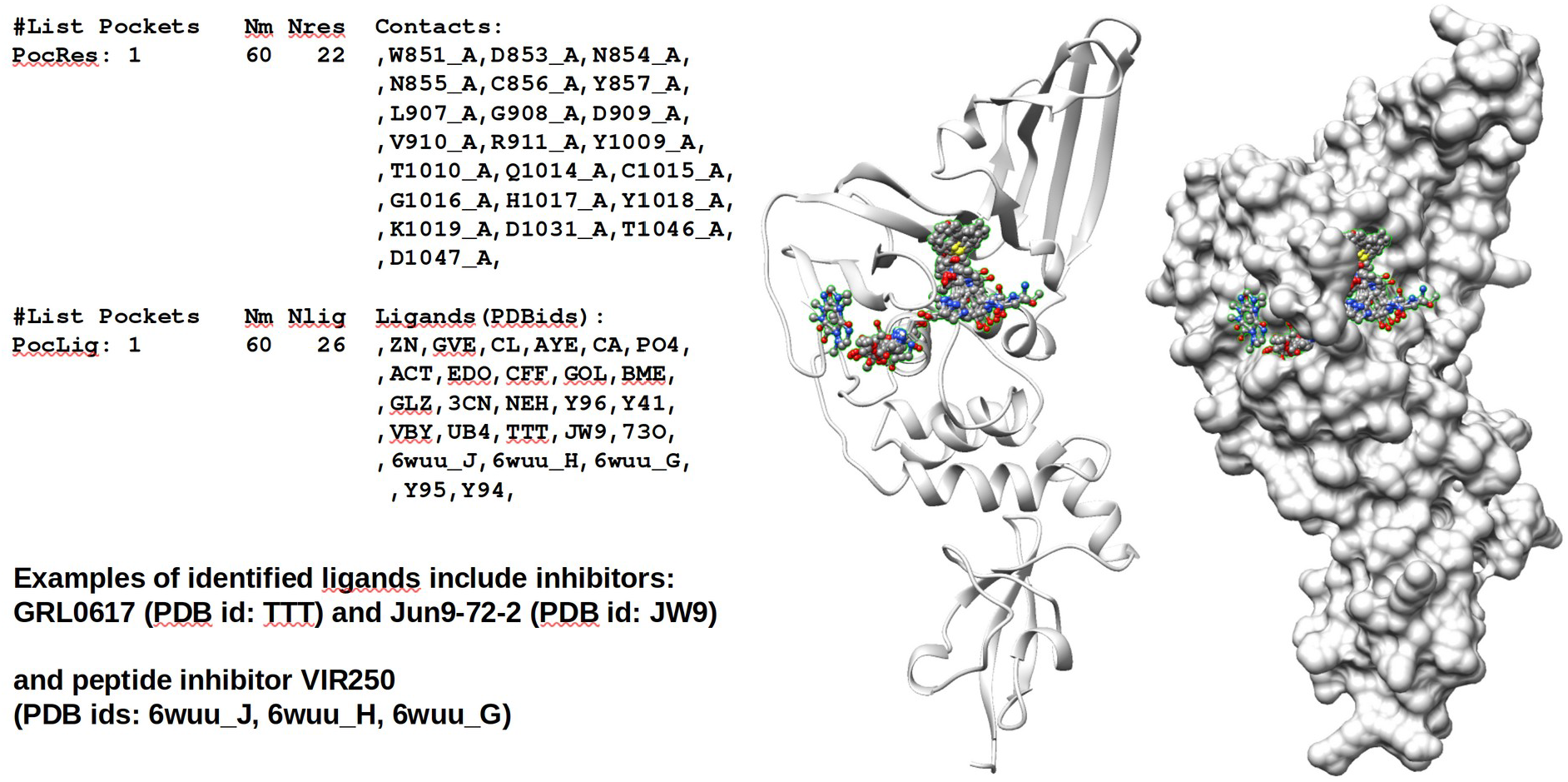
Identified in PL2pro pocket cluster #1. A list of identified ligands includes three important inhibitors: GRL0617, Jun9-72-2, and VIR250. Agreement in sets of contact residues is also observed (see Figure 4.)

The clustering of identified pocket-spheres matches and their corresponding ligands characterizes distinct pocket regions within the protein by the sets of residues interacting with inserted ligands. The clustering of interacting residues allows identification of overlapping parts of ligands that approximate the ligand’s “core” conformation within detected pockets and for each cluster defines a representative set of residues forming a consensus pocket. An example of an identified cluster of pocket-spheres is shown in Figure 2. Each protein may have more than one such consensus pocket.

### PDBspheres library

Each protein-ligand template sphere in the PDBspheres library is a subset of the records from a corresponding PDB entry, consisting of the coordinates of all atoms belonging to a given ligand, and coordinates of all protein atoms belonging to residues near that ligand (water atoms are excluded). The protein residue is part of the sphere if at least one of its atoms is within 12.0 Å of any atom of the ligand. Previous research indicated that distances of 7.5 Å are sufficient to capture informative functional properties for clustering purposes (30); however, based on our experimentation using the LGA program for detecting local structure similarities we expanded the distance to 12.0 Å. This size of the “template sphere” is sufficient to capture local structural environment of the ligand for protein structure comparative needs. Our tests indicated that larger than ∼12 Å distance spheres would affect accuracy in calculated structure conformation-based local residue-residue correspondences between template spheres and the query protein within detected pockets’ regions. On the other hand, the smaller than ∼12 Å distance criteria may not provide sufficient fold constraints to capture the uniqueness of searched pockets. To assist in identification of functional residues, PDBspheres also collects and reports information on protein-ligand interface residues. These residues are identified as those of which at least one atom is within 4.5 Å of any ligand atom.

In the PDBspheres library the 12.0 Å “sphere” entries have been constructed for each ligand in PDB, including peptides, metals, and ions, although the library includes only peptides containing 25 or fewer residues. The library is updated weekly in coordination with new PDB releases. As of 2021/10/13 the library consists of 2,069,796 spheres (binding site templates).

The primary use of the PDBspheres library is to identify “sphere” protein structures that are structurally similar to regions of a query protein structure. The query protein structure may be a complete, multimeric assembly, or it may be a single sub-unit of such an assembly, or even a fragment of a protein as long as it carries enough structural information (local structure conformation formed by atom coordinates) to reflect a structural shape of a putative binding site.

### PDBspheres similarity searches

The fundamental identification method used by PDBspheres is structural alignment of a sphere with the query protein structure, and the assessment of the structural match. For a general use, a comprehensive search – that is, a structural alignment-based search of the entire PDBspheres library (over 2 million sphere templates) – would be computationally expensive. One means to address this issue is to conduct an initial search for matches to a query protein structure on a subset of sphere templates. Then, if needed, the search can be expanded to additional templates from the library. In its current implementation for possibly faster processing, the PDBspheres library searches can be restricted to a subset of template spheres selected based on 1) ligand specificity, e.g., ligand names, their sizes or similarities, or 2) sequence similarity between the query protein and PDB proteins from which the PDBspheres library entries were derived. In the latter approach, the sequence similarity searches are conducted using the Smith-Waterman algorithm against FASTA-format sequences of all proteins contributing to the PDBspheres library entries. In the current version of PDBspheres, the Smith-Waterman algorithm “ssearch36” (31) is used.

In the PDBspheres library each sphere template entry includes a number of characteristics such as the ligand name (PDB ligand ID), the number of heavy atoms in the ligand, and the number of residues in the protein fragment forming the protein-ligand “sphere” (that is, the size of the pocket template). Thus, the list of matching spheres can be screened using specific criteria related to the expected pocket size, exact ligand name or ligand similarity. A selection of the template spheres based on the estimated similarity between the expected ligand and the PDB ligands from the PDBspheres system can be quantified by the Tanimoto or Tversky similarity indexes (32).

After a selection of template spheres is completed, each member of this set is evaluated by assessing its structure similarity with a query protein. The evaluation is performed in two steps: 1) detection of the region resembling the binding site shape, and 2) assessment of the similarity in residue conformations including a conformation in their side chain atoms. The primary search involves a calculation of structural alignment between template spheres and a query protein. The structural alignment is conducted using the LGA program and is calculated using a single point representation of aligning residues (Calpha atoms, Cbeta atoms or any other point that can represent a residue position.) The structural similarity search returns similarity scores (LGA_S) and alignments between template spheres of various pockets and the query protein. The final evaluation of identified pockets is done by assessing similarities in the conformation of all atoms (including side chain atoms) using the GDC metric.

The PDB ligand from the sphere template is translated and rotated according to the transformation that results from aligning the sphere template atoms to the query protein atoms. The ligand atoms are not considered in this alignment, and their conformation is not altered. It means that potential steric clashes between protein residues and atoms of “inserted” ligands can be observed. The number of possible clashes is reported.

PDBspheres reports the following set of characteristics for all pockets detected within a given protein:

a. PDB identifier of the protein structure from which a template sphere matching the query protein pocket was derived,
b. PDB identifier of the ligand from the template sphere matching the query protein pocket,
c. list of residues in the query protein in direct contact with the inserted ligand (residues within the distance <= 4.5 Ångstroms),
d. coordinates of the centroid of the ligand inserted into the detected pocket in the query protein.

Because the pocket searches can match template spheres to different regions of the query protein all identified pocket candidates are organized into clusters. Within the PDBspheres approach the merging and clustering of individually detected pockets (i.e., pockets detected based on individual template spheres) is performed to satisfy the following two criteria:

1. within each cluster the pockets are grouped together based on sets of query protein residues interacting with inserted ligands. When more than 80% of predicted contact residues are the same, then these sets are merged to define residues forming the pocket,
2. pockets and the sets of residues identified as in contact with ligands are merged when the inserted ligands overlap (the distances between ligand centroids are not larger than 2.0 Ångstroms.)

This grouping of pockets and their corresponding inserted ligands allows to define sets of residues interacting with similar ligands and also helps identify overlapping parts of different ligands. Such overlapping parts can approximate the ligand’s “core” conformation, whose protein contact residues can be used to define representative (consensus) pockets within the protein.

For a given protein the following set of measurements and scorings are provided to help assess the confidence level of the pocket prediction (see examples provided in Figure 4 and supplemental File1):

- LIGAND – protein-ligand template sphere identifier which includes PDB id of the ligand, ligand size, and PDB id of the protein-ligand complex from which the sphere was extracted.
- Ns - number of residues forming the protein-ligand template sphere.
- RMSD – root mean square deviation calculated on superimposed Calpha or Cbeta atoms from sphere template and detected protein pocket.
- Nc – number of conserved, i.e., “tightly” superimposed, residue pairs between the template sphere and detected pocket in an evaluated protein.
- SeqID – sequence identity in structure aligned conserved residues. The higher value indicates that the protein forming a template sphere and the query protein might be close homologs.
- LGA – structure similarity score based on aligned by the LGA program Calpha or Cbeta atoms.
- GDC – structure similarity score calculated by the LGA program assessing agreement in conformations of all atoms (i.e., including side chain atoms).
- N4 - the number of predicted protein-ligand contact residues (i.e., query protein residues observed within 4.5 Å of the inserted ligand).
- cl - the number of query protein residues that may have possible steric clashes with inserted ligand’s atoms.

Firstly, to be considered a predicted site candidate, there must be at least ten aligned residues (Nc>=10) that are conserved between the query protein and the sphere. Secondly, the structural similarity measured by GDC (29), which counts how many atoms (including side chain atoms) in the query protein and the template sphere are in close superposition, must be at least 55% (GDC>=55.0). Thirdly, there must be at least one query protein binding site residue that has an atom within less than 4.5 Å of the atom of the inserted PDB ligand (N4>=1). Finally, there are no more than two steric clashes (cl<=2) (distance less than 1.0 Å) allowed between query protein binding site residue atoms and any atom of the inserted ligand. The “cl” number is counted per residue which means that multiple clashing atoms, all within a single residue, are allowed, but not more than two residues can be involved in clashes.

We may expect a higher confidence in predicted binding sites when: Nc>=25, GDC>=65, and cl<=1. Note that these thresholds cannot be considered as absolute requirements. For example, possible clashes indicated by “cl>0” may suggest a need to correct the placement of the inserted ligand. Ideally, a score of “cl=0” would enhance the confidence in ligand-protein binding prediction, however, in many cases when “cl>0” the ligand placement within the pocket can be fixed by additional adjustment or refinement of protein side-chain atoms through docking or molecular dynamics (MD) simulations. It is also important to keep in mind that the query protein structure may not be in its “holo” conformation to properly accommodate the ligand (i.e., without clashes), so the conformation of binding site residues of the protein may require more substantial optimization. Indeed, recent studies of ligand binding site refinements show significant success in generating reliable “holo” (ligand-bound) protein structures from their “apo” (ligand-free) conformations (33).

If N4 – the number of protein atoms within 4.5 Å of the ligand – is very low, then this indicates that the current placement of the ligand does not show strong interactions with residues of the protein binding site. This may suggest that the pocket is too large (or some residues in the protein model are missing, or side-chain atoms are not in the right conformation). Of course, it may also indicate that the identified pocket is incorrect, or the location of the inserted ligand is wrong, but these conclusions can only be confirmed by more detailed inspection. In such cases it can be informative to check the overlap of the template sphere pocket and the query protein pocket (i.e., check the Nc number of conserved superimposed pocket-forming residues). Higher overlaps (e.g., Nc>=25 or more) indicate greater size of the region forming the pocket and thus the higher confidence in reported pocket similarities (more residues identified as conserved). However, such thresholds cannot be decided upfront because in cases of shallow cavities or interface sites on the surface of the protein the Nc number can be low.

The application of the PDBspheres for binding site detection in papain-like proteinase (PL2pro) from COVID-19 is illustrated in Figures 2-4. In Figure 2, we show results from detected pocket cluster #1 which was defined based on PL2pro pocket structure similarities with 60 pocket-ligand template spheres (Nm=60). The number of predicted different bound ligands is Nlig=26, and the combined total number of residues being in contact with bound ligands is Nres=22.

In Figure 3 we show that the poses of three inhibitors from pocket cluster #1: VIR250, GRL0617, and Jun9-72-2, whose PDB ids are: 6wuu_J, TTT, and JW9 respectively, overlap significantly. These ligand poses are brought from 11 different template spheres listed in Figure 4: three for VIR250, five for GRL0617, and three for Jun9-72-2.

**Figure 3.**
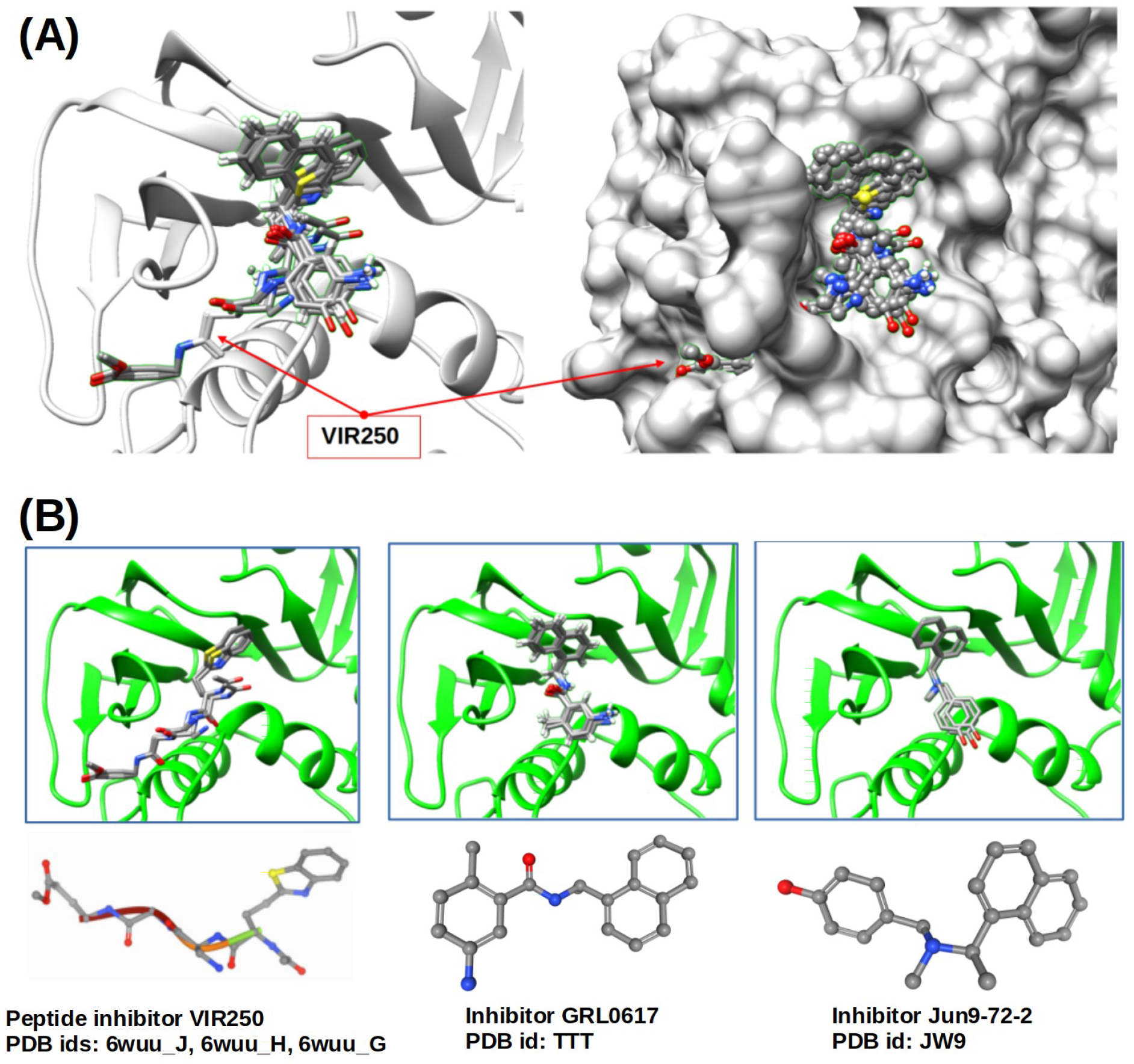
(A) Example of the peptide inhibitor VIR250 that passes through the “gorge”. Inhibitors GRL0617 and Jun9-72-2 are placed only on one side (right part) of the cavity of pocket cluster #1. (B) Assessed by PDBspheres poses of all three ligands that come from different template spheres show strong overlap.

**Figure.4.**
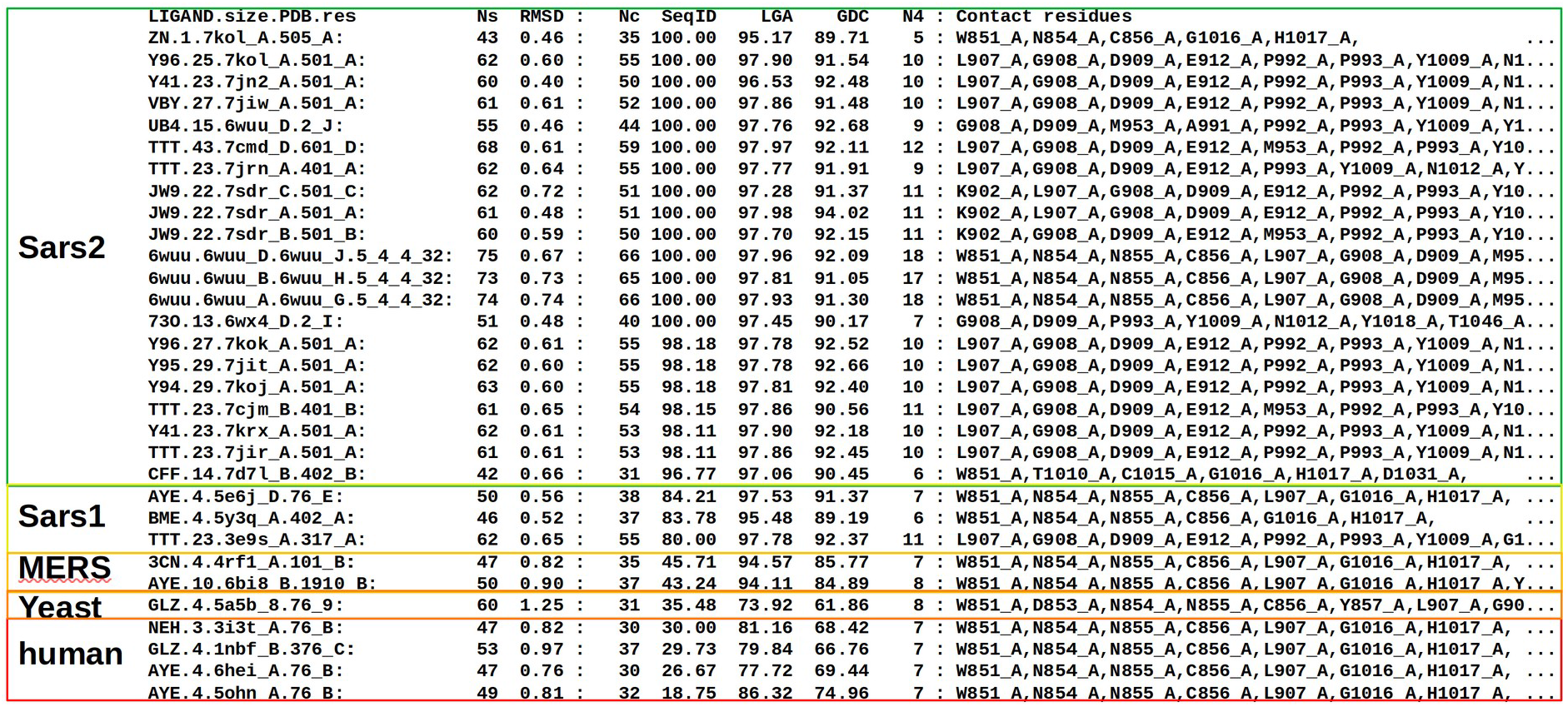
Fragment of the PDBspheres summary table reporting predicted PL2pro ligands assigned to pocket cluster #1. Results highlighted in green (when SeqID “>95”) indicate that similar pockets are detected in Sars2 or variants of Sars2. When SeqID “<30” (results highlighted in red) examples of similar pockets detected in human proteins are shown. An agreement in sets of contact residues indicates that all listed ligands are predicted to bind the same pocket identified through clustering as “pocket cluster #1”.

In Figure 4 we present a snapshot (only 31 template spheres out of Nm=60) from the summary table automatically created by the PDBspheres system for pocket cluster #1. A complete summary table of predicted pocket-ligands for a structural model of papain-like proteinase (PL2pro model: nCoV_nsp3.6w9c_A.pdb) is provided in supplemental File1.

### Benchmark dataset and evaluation metrics

The benchmark dataset used to assess the performance of PDBspheres is the “LBSp dataset” described in Clark et al. (25, 26). It comprises 304 unique protein families (2,528 structures in 1456 “holo” and 1082 “apo” conformations) with each family represented by several protein structures in different possible conformations. Since this LBSp dataset is curated, well designed, large and diverse, we have found it very suitable to test the performance of the PDBspheres method comprehensively.

The main criterion often used for the assessment of the accuracy of predictions of the binding site is the agreement in the set of residues predicted to be in contact with experimentally confirmed ligands (residue contacts derived from protein-ligand co-crystals from PDB (27)). To estimate the accuracy of our method we followed the same metrics and evaluation procedures as used in (25). The authors defined reference lists called unified binding sites (UBS) as unions of all residues contacted by any bound ligands within the protein family. The scoring of binding site predictions is determined by agreement with the UBS reference using calculated numbers of any residue predicted to be part of the binding site which is confirmed in the experimental data is denoted as true positives (TP), all remaining residues (which are not predicted as part of the binding site and not confirmed in experimental data) are denoted as true negative (TN), any residue predicted to be part of the binding site which is not confirmed in experimental data is denoted false positive (FP) or over-prediction, and any remaining residues in the experimental data not accounted for in an algorithm’s predicted binding site are denoted as false negative (FN) or under-predictions.

Both Matthew’s Correlation Coefficients (MCCs), and F scores as calculated by formulas (Eqs. 1-2) have been used as metrics to represent the predictive power of evaluated binding site prediction methods.

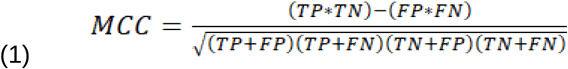

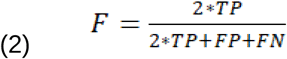

Of course, there are other metrics sometimes used to evaluate protein-ligand binding sites detection successes, e.g., distance center center (DCC) (34) and discretized volume overlap (DVO) (12). DCC is a metric to evaluate the correct location of the pocket based on the distance between the predicted and the actual center of the pocket (centroids). If DCC is below 4 Å, the pocket is considered as correctly located. DVO, on the other hand, is defined as the ratio of the volume of the intersection of the predicted and the actual segmentations to the volume of their union. It assesses the correctness of the shape of predicted pockets. However, in our study we follow metrics described in Clark et al. (25) as they focus on assessing accuracy of binding site prediction methods based on the correctness of identified pocket residues being in contact with ligands. Similar approaches were used in CASP experiments to evaluate the performance of binding site prediction methods.

## RESULTS AND DISCUSSION

### Performance of PDBspheres on LBSp dataset

In Clark et al. (25), predictive power of different methods is assessed using two metrics: F scores and Matthew’s Correlation Coefficients (MCCs). The F scores and MCCs provide a good description of the relative success of evaluated algorithms by assessing whether or not a method produced a predicted binding site that contains residues in common with the reference UBS list of ligand contact residues. They assign high scores not just when at least one residue in a given site is identified correctly, but rather reward methods with more correct and less false predictions of contact residues implying which of the algorithm pocket predictions are close to the “correct” location on the binding surface of the protein. Note that a similar evaluation procedure using MCC scores between predicted and observed contact residues was also applied in CASP experiments. Clark et al. (25) compare the following methods on their LBSp dataset: Surfnet (19), Ghecom (35), LIGSITE (36), Fpocket (37), Depth (38), AutoSite (39) and Kalasanty (15). Five of these methods are considered to be geometry-based, while one is energy-based, and one is machine-learning-based. To perform our analysis, we took prediction results for these seven methods from the supplemental data from the publication (25). In Table 1 we present the prediction results for PDBspheres and recalculated – for consistency - F and MCC scores of the seven mentioned above methods. The last six data rows in Table 1 show results from the evaluation of PDBspheres when the pocket detection was performed using the complete PDBspheres library (100%), and when the PDBspheres library was restricted to templates with no more than 90%, 80%, 70%, 60% and 50% sequence identity with query proteins, respectively. The six rows for PDBspheres include also the mean values of MCC in addition to the median values since this metric was used by CASP assessors.

### Structural similarity of binding sites vs. sequence identity

Here, we discuss how close in sequence identity two proteins need to be to have structurally similar binding sites, addressing these two questions:

- How low the level of sequence identity between two proteins can be and still share similar pockets and perform similar functions?
- Do such proteins have enough similarity in their functional sites to bind ligands in a similar manner?

Figure 5 illustrates results from the LBSp database pocket identification calculations, which were summarized in Table 1. These results illustrate that in contrast to the large diversity of possible protein sequences the number of structurally distinct pockets is limited, therefore proteins that by sequence are very different may still share almost identical structural conformations in local regions (e.g., binding sites) and can perform similar functions. This observation served as a basis for the development of our PDBspheres structure template-based binding site detection method. The only limitation in its ability to identify correct pockets for a given protein might be an underrepresentation of particular pocket’s conformation in currently experimentally solved protein structures deposited in the PDB database. For example, in case when a non-restricted library is used the binding sites for all 304 families (“apo” and “holo” protein conformations) were predicted. However, in the case of the restricted library (which excludes template spheres derived from proteins showing more than 90% sequence identity to the query proteins) the binding sites for 5 families of “apo” versions and 7 families of “holo” versions were not predicted (see Table 2). In Table 2 we show even more results indicating that the protein binding sites are highly structurally conserved and can be successfully detected using structure-based template spheres taken from proteins sharing low sequence identity with targeted proteins. Even when the restrictions applied to the PDBspheres library increase from 90% to 50% sequence identity, the success in detecting the number of predicted pockets does not drop so rapidly.

**Table 2.** Number of detected pockets and pocket’s families when the PDBspheres libraries are restricted to templates with sequence identity to proteins from the LBSp dataset not exceeding introduced cutoffs. Results in this table reflect correctness in detection of just the pocket location in the protein, not an assessment of the accuracy and completeness of predicted interacting residues which are reported in Table 1.

| Restricted sequence identity | Number of detected pockets |  |  | Number of pocket's families |  |  |
| --- | --- | --- | --- | --- | --- | --- |
|  | All | Apo | Holo | All | Apo | Holo |
| 100% | 2528 | 1082 | 1456 | 304 | 304 | 304 |
| 90% | 2481 | 1056 | 1435 | 299 | 299 | 297 |
| 80% | 2469 | 1051 | 1428 | 298 | 296 | 297 |
| 70% | 2469 | 1051 | 1428 | 298 | 296 | 297 |
| 60% | 2464 | 1048 | 1426 | 297 | 295 | 296 |
| 50% | 2457 | 1046 | 1421 | 295 | 294 | 294 |
|  | Percent of detected pockets |  |  | Percent of pocket's families |  |  |
| 50% | 97.2% | 96.7% | 97.6% | 97.0% | 96.7% | 96.7% |

**Figure 5.**
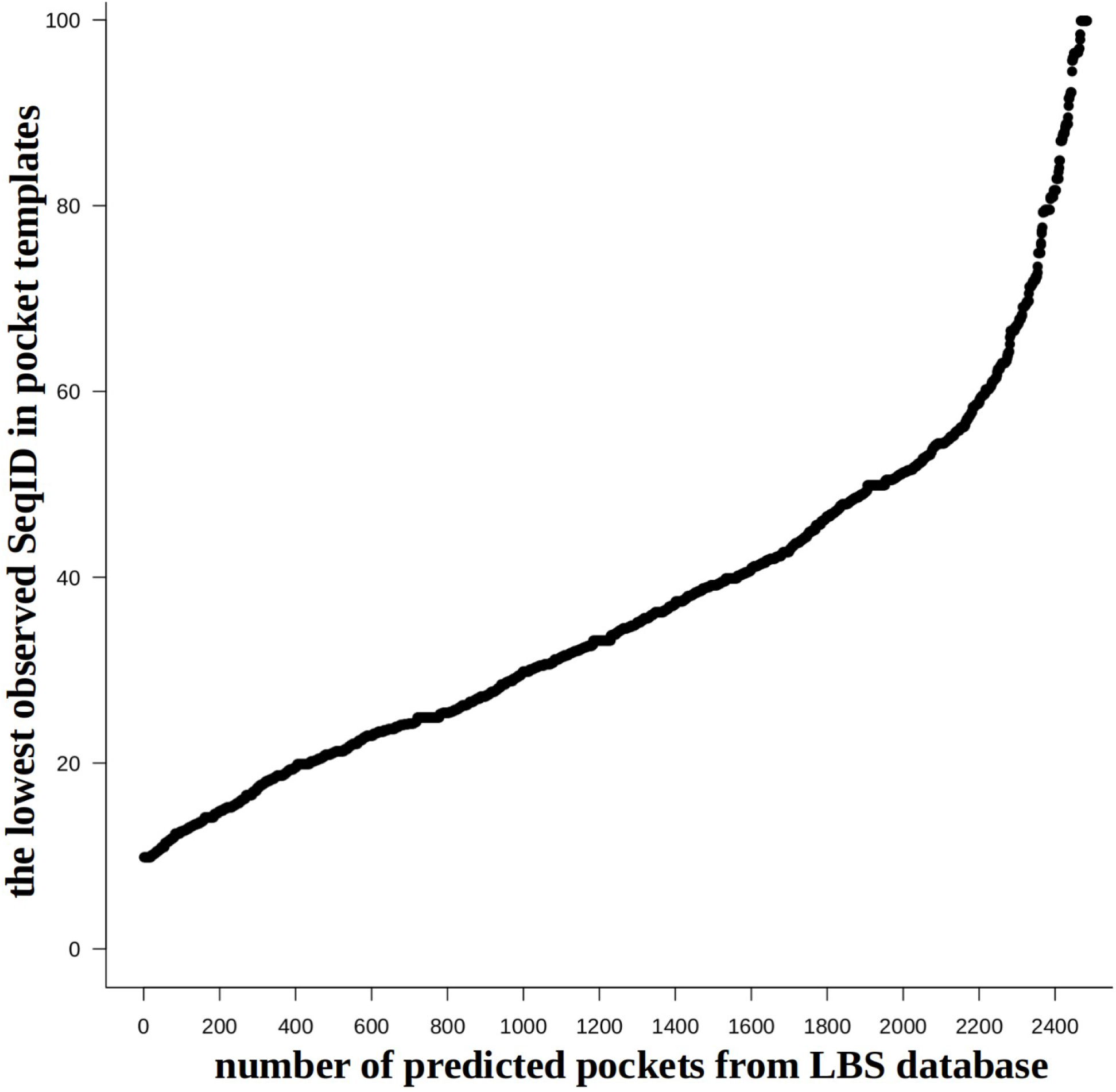
The plot illustrates how many pockets from the LBSp database proteins (all 2,528 pockets; “apo” and “holo” combined) can be predicted based on binding site templates derived from proteins with lowest possible sequence identity to the query proteins. For example, at least 400 pockets (see left bottom part of the plot) can be predicted based on template binding sites derived from proteins that share as low as 20% sequence identity (SeqID) with a query protein. On the other hand, more than 2,400 pockets (∼95%) can be correctly predicted based on templates sharing no more than 80% of SeqID with query proteins in their binding site regions.

The same can be observed in Figure 5. More rapid decline in the ability to successfully detected pockets starts when the restriction on the library gets below 50% of sequence identity. Let us note that the main difference between results shown in the Table 2 and Figure 5 is that the plot illustrates the correctness of the prediction (i.e., as accurate as possible identification of residues involved in protein-ligand interaction) while in Table 2 we report results from just detection of the pocket location in the protein (i.e., without any assessment of the accuracy and completeness of predicted interacting residues).

The PDBspheres method leverages this structure conservation quality efficiently. Indeed, thanks to the richness in diversity of protein structures deposited in PDB the system is able to identify location of 97% binding sites from the LBSp dataset using structural templates that share as low as 50% sequence identity with targeted proteins. And even with such significant restriction to the library of template spheres the accuracy in identified sets of residues interacting with ligands is still high ∼0.75 (median) and ∼0.7 (mean) as measured by F and MCC metrics (see Table 1).

As mentioned in Clark et al. (25), we expected that the template-based method such as PDBspheres could outperform non-template-based methods which is evidenced in Table 1. We should emphasize, however, that PDBspheres is an exclusively structure-based method which does not utilize any prior information from libraries of sequences/residues forming binding sites. Additionally, PDBspheres does not use any prior information about the location of searched pockets in proteins from the same family. We treat all structures equally and independently (as pairwise comparisons) in finding structural similarities between the query protein and template spheres. Of course, we can find such cavities more easily in the “holo” conformations, but as the results show, we can also find adequate structure similarities in “apo” versions of the query protein; again, strictly based on similarities in structure without any sequence-based knowledge of residues forming the pocket (residue information that could be transferred from some databases of “holo” structures.) All results reported here are based on PDBspheres superpositions calculated on C-alpha atoms. Results based on superposition of C-beta atoms (results not shown) are virtually identical.

In Figure 6 we show an example of correct pocket detection by PDBspheres using different pocket-ligand template spheres derived from proteins that share low sequence identity. Two compared proteins (serine proteases) have less than 38% of sequence identity, but they are structurally similar at the level of over 86% by LGA_S (assessment of similarity on the C-alpha atoms level) and 77% by GDC (all atoms level). Of the residues that are in contact with corresponding ligands (distance below 4.5 Å) in the identified similar pockets, 11 out of 17 are identical (65%). The pocket template spheres come from two PDB complexes that bind inhibitor compounds having PDB ligand ids 2A4 and QWE, respectively. These ligands are significantly different in size. Therefore, the distance between ligand centroids calculated from the complexes of the same orientations is very different, specifically, the distance between centroids of ligands when inserted in any of these pockets is about 6.0 Å. However, the portions of the ligands inserted in pockets have a similar overlapping region and are in contact with similar amino acids from the pockets. In each of these two protein-ligand complexes the core parts of the ligands are in contact with 13 residues of which 11 are identical (85%) (see Figure 6 (C) and (E)). We have had a similar observation in the example of Human Neuropilin-1 illustrated in Figure 1, when we focused on possible discrepancies in distances between centroids of two experimentally confirmed orientations of the same ligand inserted into the same pocket. The distance between “core” parts of the ligand was very small, while the outside parts varied significantly. It suggests that if we are interested in identifying protein residues critical for ligand binding, we should concentrate mostly on those residues that are in contact with ligand’s “core” parts. Likewise, as shown in Figure 6, in case of different proteins and different ligands a high sequence identity of residues being in contact with “core” parts of the ligands can help identify those residues that may be critical for binding in similar pockets. These results illustrate how PDBspheres can be used to detect conservation in local structural conformations and to assess conservation of critical contact residues; both can assist in inferring protein function. These conclusions are also supported by results from evaluation of large and diverse set of proteins collected in the PDBbind dataset (40).

### Clustering binding sites from PDBbind database

In this section, we describe how PDBspheres can be used to perform structure-based clustering and structure similarity analysis of binding sites from the PDBbind dataset (an extensive collection of experimentally measured binding affinity data for the protein-ligand complexes deposited in PDB) (40) (ver. 2019). Some of these results were leveraged in our previous work generating rigorous training and validation datasets for machine learning of ligand-protein interactions (13). Here we want to address the following questions:

- To what extent can structurally similar binding pockets having similar ligand placement allow inference of binding affinity from one pocket-ligand pair to another pocket-ligand pair?
- To what extent can clustering of detected pockets and calculated structure similarities among clustered pockets from different proteins provide functional information for protein annotation?

In our analysis, we focus on the “refined” 2019 dataset (4,852 structures) which we expanded by adding 24 structures from the previous PDBbind release that are not present in the 2019 version. Hence, in total the dataset of evaluated binding sites consists of 4,876 structures. Since we are interested in the assessment of similarities between specific pockets listed in proteins from PDBbind (the PDBbind database reports only one pocket for each protein regardless of how many different pockets a given protein may have or how many alternative locations of a given pocket in a multichain protein complex can be observed), we restricted our structure similarity searches and evaluations to only those regions in PDBbind proteins that encompass targeted pockets. Note that many protein structures in the PDBbind refined dataset are multidomain complexes with total sizes of more than 2000 residues, so they can have multiple binding sites in addition to the targeted ones. Therefore, in our approach to evaluate similarities and cluster pockets from PDBbind, each of its protein structures was reduced to the region in the close vicinity of a reported ligand (residues having any atom within 16 Å of any ligand atom), and each ligand binding site reported in PDBbind was associated with its corresponding PDBspheres template sphere (residues having an atom within 12 Å of a ligand atom). We performed an all-against-all PDBspheres detection and pocket similarity evaluation using the 16 Å protein region representation of each PDBbind protein. Results from the pairwise pocket similarity evaluation are provided in supplemental File5. Structure similarity results allowed grouping of the 4,876 PDBbind protein pockets into 760 clusters. Cluster details are provided in supplemental File2, File3, and File4. Figure 7 illustrates the clusters, and it is a snapshot taken from the HTML supplemental File6 (interactive overview of predicted clusters created using “plotly” R graphic library). Each axis of Figure 7 indexes a sample of 4,876 protein pockets, where sampled protein names are reported to help identification of corresponding pockets or clusters on the plot. A “zoom-in” option in the “plotly” graph expands each rectangle to show a list of individual members of a selected cluster. It allows for each of the sampled proteins to be labeled with its exact location within the cluster which is shown as the example in Figure 8. Further illustrations can be found in supplemental File6. Each rectangle in Figure 7 represents a cluster and is composed of small markers – one for each sampled protein-pocket pair within the cluster – colored by the GDC all-atom similarity score between each member of the cluster, where colors closer to red indicate a higher degree of similarity between members. For example, the first three large rectangles at the bottom left of Figure 7 represent predicted clusters: cluster #22 (318 members), #8 (351 members), and #5 (322 members), respectively. Evaluation of predicted clusters which is reported in supplemental File2 shows that all members of cluster #22 are enzymes which belong to EC subclass 4.2.1.1, members of cluster #8 - EC 3.4.21, and cluster #5 - EC 3.4.23. All clusters along the diagonal in Figure 7 have high within-cluster similarity and are separated from other clusters according to the applied “exclusive” clustering approach. Some of automatically created by PDBspheres clusters are formed by grouping together proteins from several “finer” subclusters. For example, at the top right there is a large cluster (#25 with 382 members assigned to several subclasses of the broad-spectrum transferases - EC class 2.7 (“Transferring phosphorus-containing groups”), with varying degrees of similarity among its members as they belong to different and more specific subclasses which are still similar enough (according to the selected thresholds) to form one distinct cluster #25. Data used in Figure 7 are reported in supplemental File2 and File6.

**Figure 6.**
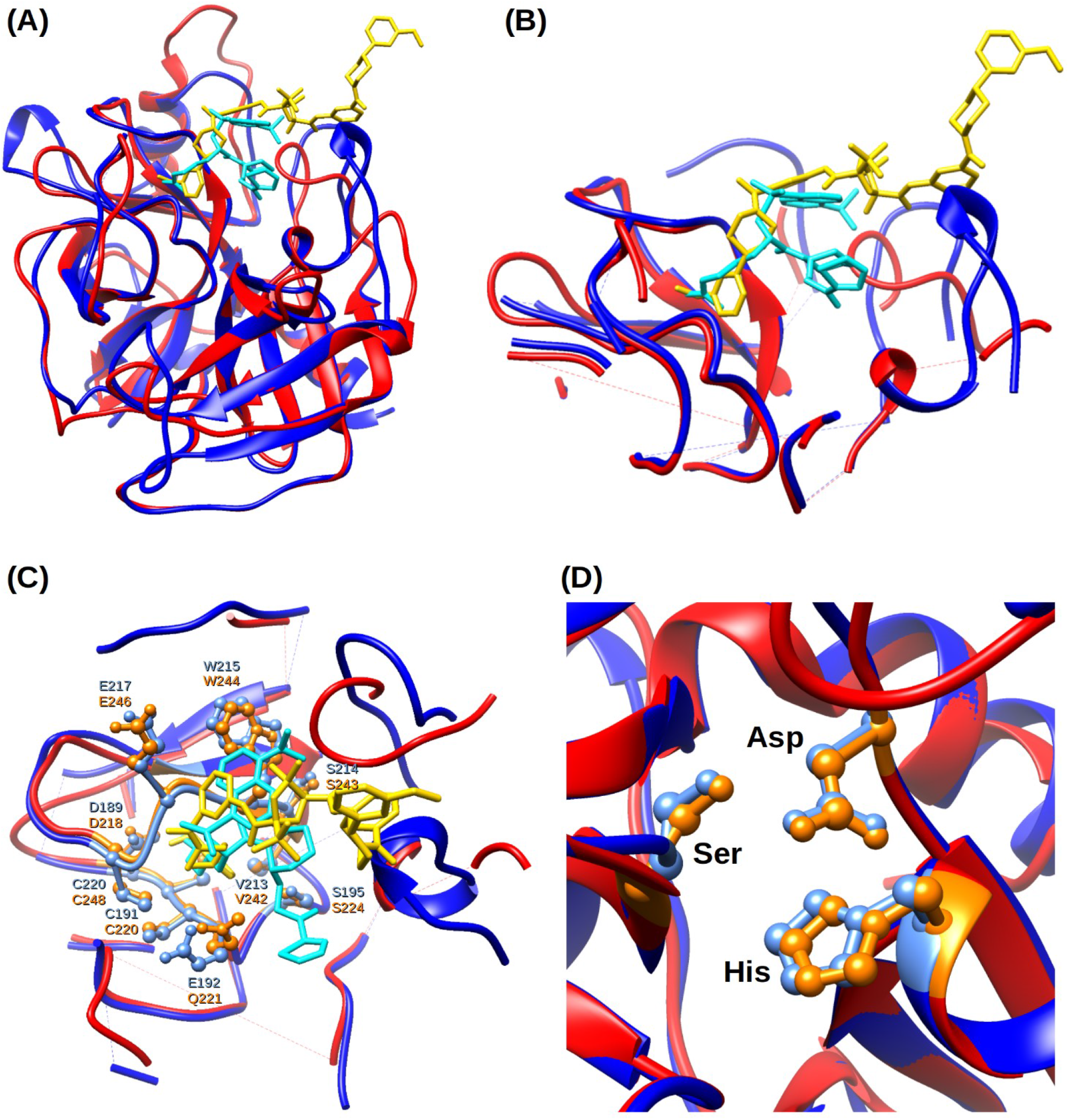
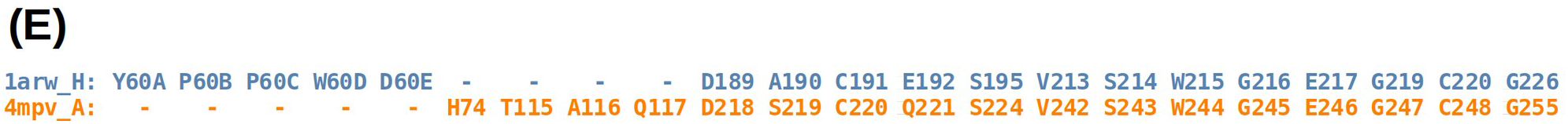
Example of two structures of serine proteases: Tryptase (in red PDB chain: 4mpv_A) and Thrombin (in blue PDB chain: 1a4w_H) that share high structure similarity in their binding sites (over 77% by GDC) while the level of sequence identity between them is no higher than 38%. (A) Overall structure superposition of two protein-ligand complexes showing location of bound inhibitors. (B) PDBspheres-based local superposition of corresponding protein spheres surrounding ligands. (C) Structurally superimposed spheres of 4mpv and 1a4w show significant similarity in side-chain conformation of residues interacting with corresponding ligands 2A4 and QWE. Residues interacting with ligands 2A4 and QWE are highlighted in orange and light blue, respectively. (D) Local superposition indicates a perfect agreement in the nearby catalytic triad residue conformations (His, Asp, Ser). (E) Structural alignment of residues from close distance (4.5 Å) from the corresponding ligands shows that 11 out of 13 (85%) of residues that are in contact with similar core parts of the ligands are identical.

**Figure 7.**
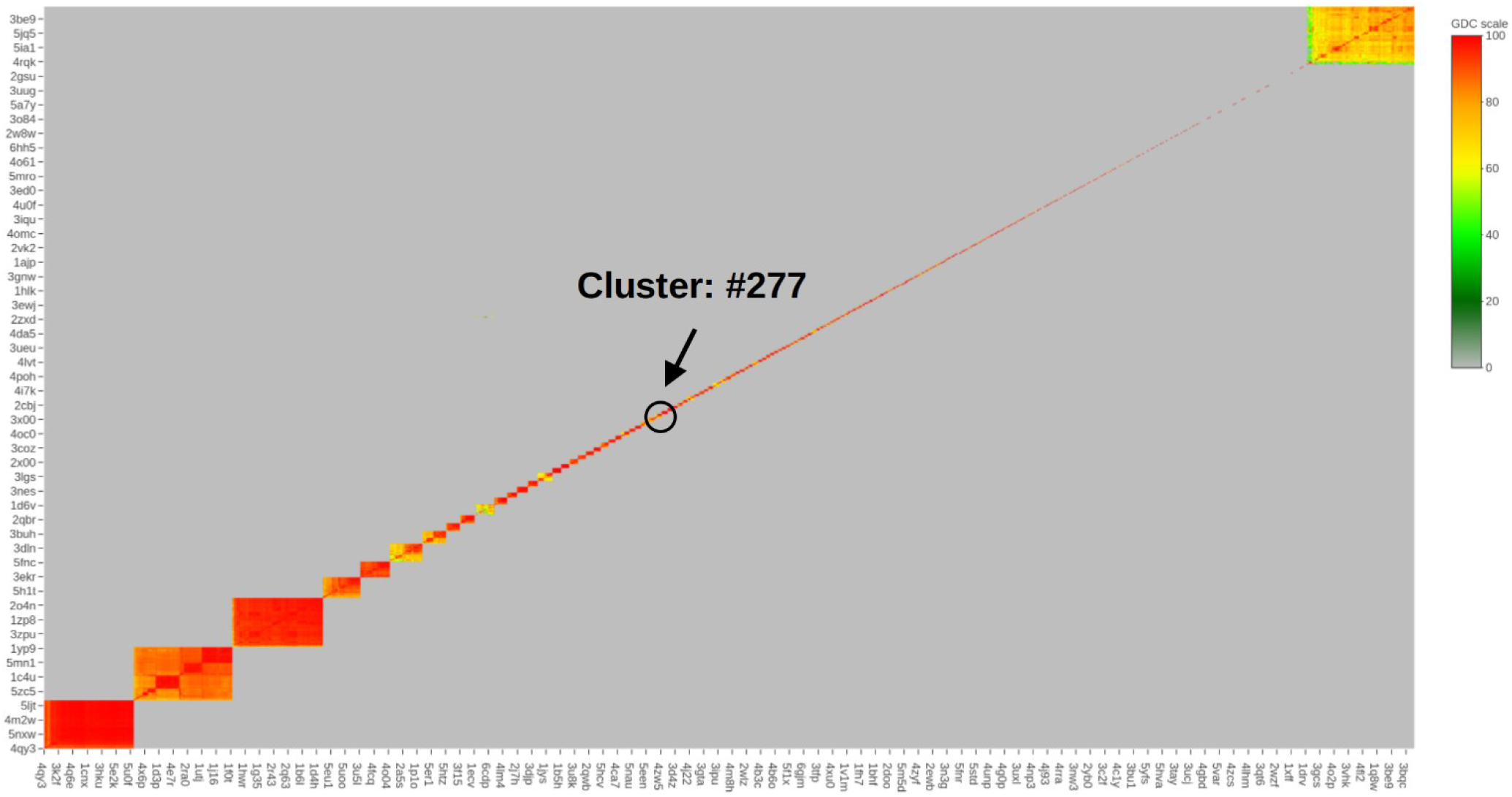
Clustering of the refined set of 4,876 pockets from the PDBbind based on their structure similarity. PDBspheres-based pocket detection and similarity evaluation resulted in 760 constructed clusters. HTML file allowing interactive overview of predicted clusters is provided in supplemental File6. Figure 8 shows a “zoom-in” to cluster #277, which contains 24 protein-ligand pairs.

**Figure 8.**
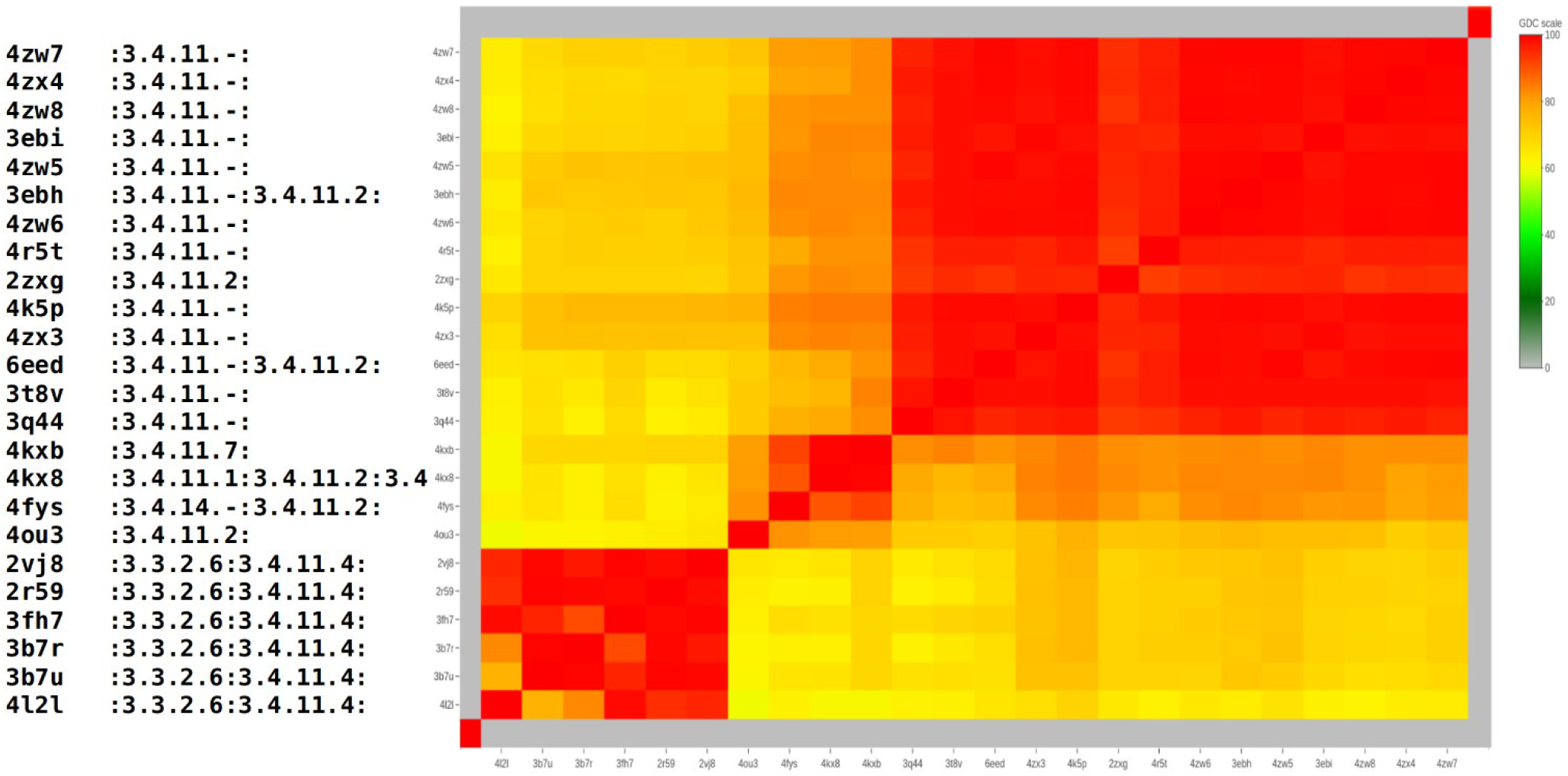
“Zoom-in” to cluster #277 identified by PDBspheres in PDBbind dataset (see Figure 7 and supplemental File2). All 24 proteins within this cluster belong to the same enzyme subclass 3.4. The first column provides PDB ids of each enzyme from the cluster. Corresponding EC numbers are reported in the second column, where alternative EC assignments are separated by colon”:”. Coloring on the plot reflects pairwise similarity scores between clustered proteins as measured by GDC.

As an example of results from PDBspheres clustering, Figure 8 shows a “zoom-in” to cluster #277 (circled and marked by an arrow in Figure 7), which contains 24 protein-ligand pairs. This “zoom-in” shows details of PDBbind binding sites grouped together based on all-against-all structure similarity measured by the GDC score. Each of the proteins grouped into this cluster share the same EC subclass (3.4) and GO annotation (:0006508:0008237:0008270:), which indicate that the PDBspheres-based clustering can assist in making predictions useful in protein functional annotation. The details of cluster #277 are reported in supplemental File2 and File4. As shown on the plot, all 24 proteins belong to the same enzyme subclass 3.4. - hydrolases that act on peptide bonds, specifically hydrolases with EC 3.4.11 cleave off the N-terminal amino acid from a polypeptide. In addition to the general functional clustering of proteins, the PDBspheres clustering approach provides finer subclustering of proteins within predicted clusters. The bottom 6 proteins from cluster #277 form a clear subcluster as they share additional EC 3.3.2.6 - bifunctional zinc metalloprotease activities.

In supplemental File2, File3, and File4 we provide specifics of assigned EC, SCOP and GO annotations, respectively, for each member of created clusters. The first two columns in Figure 8 are just a snapshot of what kind of information is reported in these files. Closer examination of these assignments for members of each cluster shows that similar pockets that are clustered together by PDBspheres share similar functions.

Another important question is: can we transfer binding affinity scores from one ligand binding site to another if the pockets and the ligand placements within the pockets are similar? Results from PDBspheres analysis suggest that pockets from PDBbind that share high structure similarity and have similar ligand placements (distances between the centroids of inserted ligands are no greater than 0.5 Å) show also similarities in reported protein-ligand binding affinities. In particular, for PDBbind entries with known Kd values, in Figure 9 (A) we compare the Kd values of each pair of 1,544 protein-ligand complexes when GDC is greater than 95% and aligned ligand centroids are within 0.5 Å. The R^2^ and Spearman values of 0.5 and 0.7, respectively, indicate the strength of the relation. Although reported scores are not very high, they can still suggest that with required similarity constraints the similar protein-ligand pairs tend to have similar Kd values. Similarly, in Figure 9 (B), the pockets with given Ki values and the same similarity constraints show the R^2^ and Spearman values of 0.44 and 0.574 for 5,137 evaluated protein-ligand pairs.

**Figure 9.**
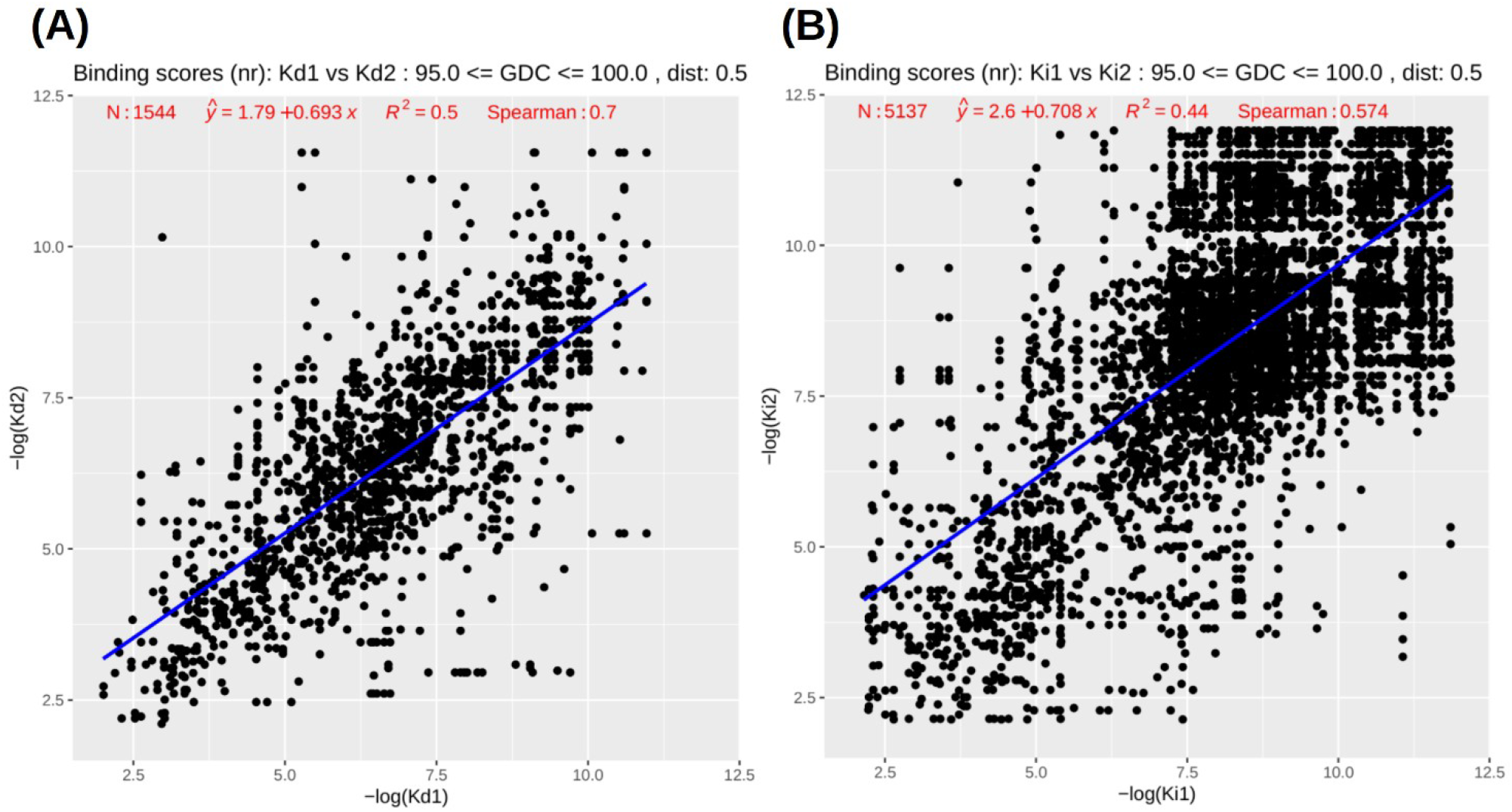
(A) Scatter plot of Kd values from 1,544 pocket pairs, and (B) Ki values from 5,137 pocket pairs. Results shown on the plot were calculated for pairs satisfying the following similarity constraints: pocket GDC similarities >=95%, and ligand centroid’s distance cutoff of 0.5 Å. Redundant pairs – self-comparisons and symmetry duplicates – are removed from calculations.

These similarities in binding affinities are even higher between pockets from different proteins when they bind the same ligand. The corresponding R^2^ and Spearman scores for 194 pairs with Kd values are 0.5 and 0.738, and for 411 pairs with Ki values are 0.69 and 0.772, respectively. Even if we relax the similarity constraints by requiring a ligand centroid’s distance cutoff of 1.0 Å and GDC pocket similarities as low as 90%, for pocket pairs that bind the same ligands, as shown in Figures 10 (A) and (B), for the Kd subset of 287 pairs the R^2^ and Spearman values are 0.46 and 0.703, and for the Ki subset of 631 pairs the scores are 0.64 and 0.752, respectively.

**Figure 10.**
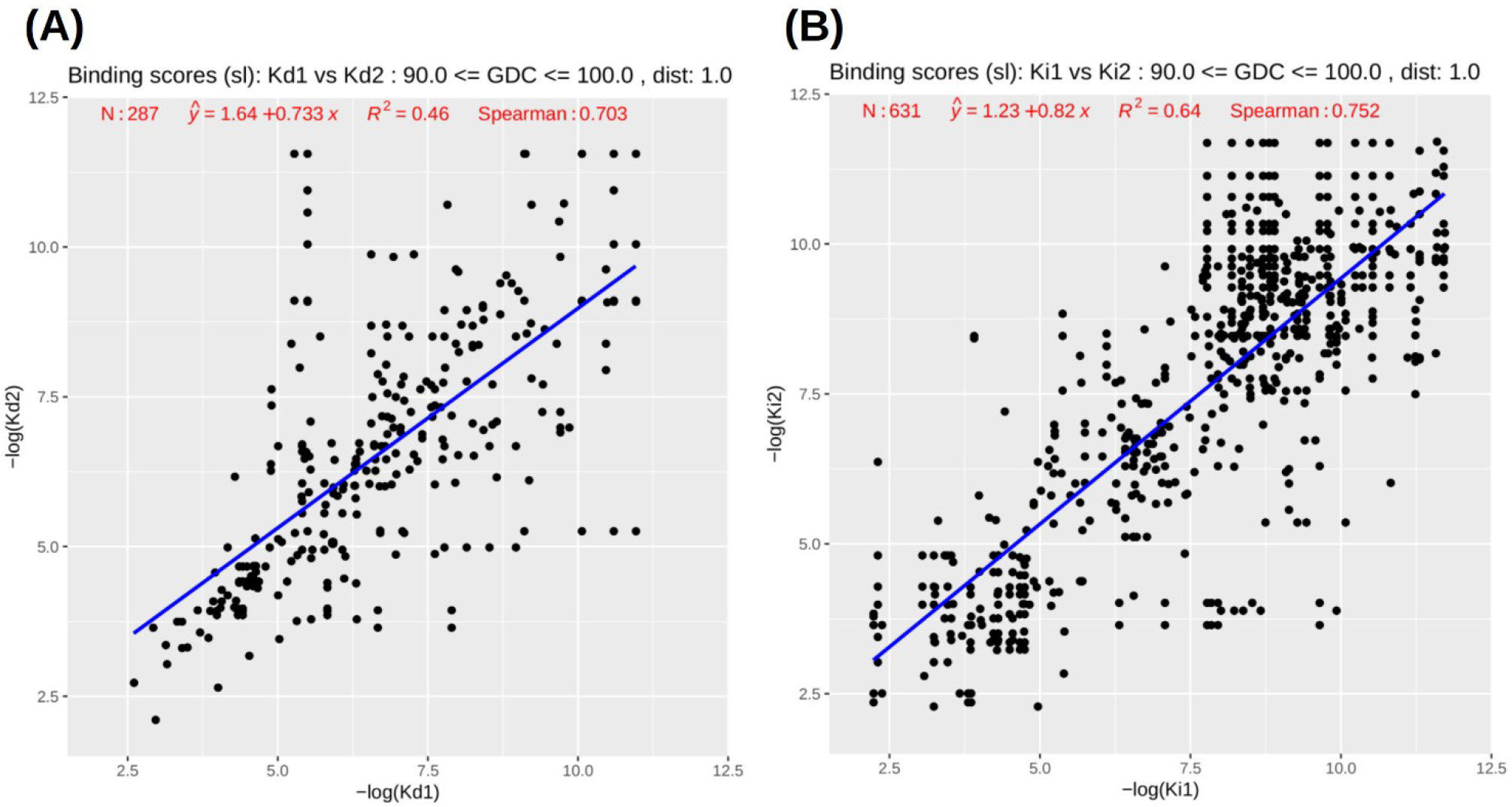
(A) Scatter plot of Kd values from 287 “same ligand” pocket pairs, and (B) Ki values from 631 “same ligand” pocket pairs. Results shown on the plot were calculated for pairs satisfying slightly relaxed similarity constraints: pocket GDC similarities >=90%, and ligand centroid’s distance cutoff of 1.0 Å. Redundant pairs – self-comparisons and symmetry duplicates – are removed from calculations.

Results from the PDBspheres clustering of the PDBbind dataset suggest that the structure similarity between pockets/ligand placements is a significant characteristic that can allow prediction of similar binding affinity values. In future work, we anticipate enhancing current measurements adding information about the specific atom location of protein residues interacting with the ligand, which may improve our predictions. Complete results from PDBspheres analysis of pairwise similarities in binding sites between proteins from PDBbind are provided in supplemental File5.

## CONCLUSIONS

While developing PDBspheres we focused on two goals: 1) binding pocket detection; and 2) identification of characteristics and scores to assess similarities between pockets to help further protein functional characterization and clustering.

In particular, with regard to binding pocket detection we find that PDBspheres’ strictly structure-based approach can correctly predict binding site regions in protein structures known to be in a “holo” (i.e., ligand-binding) conformation as well as in protein structures in “apo” (without a ligand present) conformation. While we are not directly comparing our method with other template-based binding site prediction methods we can estimate that PDBspheres has very similar accuracy when assessed by Matthew’s correlation coefficient (MCC) between predicted and observed binding residues. CASP experiments (CASP8, 9 and 10) indicated that in comparison with other methods the template-based methods were performing on the top with reported average MCC scores slightly above 0.7. For example, the average scores for the best performing template-based methods at CASP10 Firestar and SP-ALIGN were 0.715 and 0.707, respectively. Results from testing PDBspheres on LBSp dataset, reported in Table 1, show that the average MCC score for PDBspheres is at the similar level ∼0.71 when the closest to the targeted protein homologous template spheres are being removed from the library. As shown in Table 1, when the sequence identity drops, there is an immediate decline in the average MCC values from ∼0.8 (for 100% SeqID when the exact or very close match can be found in the library) to about 0.7 (for 90% to 50% SeqID when possible best matches are being eliminated). Interestingly, in case of restricted libraries the average MCC value seems to reach a steady level of ∼0.7 which agrees with results of evaluation of template-based methods in CASP8 to 10 experiments (22-24).

Since local regions in functionally similar proteins are remarkably conserved in their structural conformations, the PDBspheres method allows detection of similar binding sites even in proteins that share very low sequence similarity. Based on this observation and our tests on the LBSp benchmark dataset with restricted libraries, we can expect that structure-template-based methods may successfully predict 96% of pocket locations and reach the accuracy level in predicted binding residues measured by the average MCC of ∼0.7 for proteins that have no close representation in used libraries.

With regard to characterizing and evaluating binding pocket similarities among proteins we find that a high level of sequence similarity between different proteins is not essential to identify structurally similar binding sites and that proteins even significantly different by sequence may perform similar functions. In the structure-based detection of binding sites the similarity assessed based on calculated structural alignment using C-alpha atom positions is sufficient, and the use of other residues (e.g., C-beta atoms, or other points representing residue) in the similarity assessment does not yield better results. Structurally similar binding pockets having similar ligand placements allow inference of binding affinity from one pocket-ligand pair to another pocket-ligand pair. PDBspheres-based clustering of detected pockets and calculated structure similarities among pockets from different proteins provide information that can significantly help protein function annotation efforts.

## AVAILABILITY

The PDBspheres library and the LGA program are made publicly available for download at http://proteinmodel.org/AS2TS/PDBspheres and http://proteinmodel.org/AS2TS/LGA

## SUPPLEMENTARY DATA

Supplementary Data are available at NAR online.

**File1:**
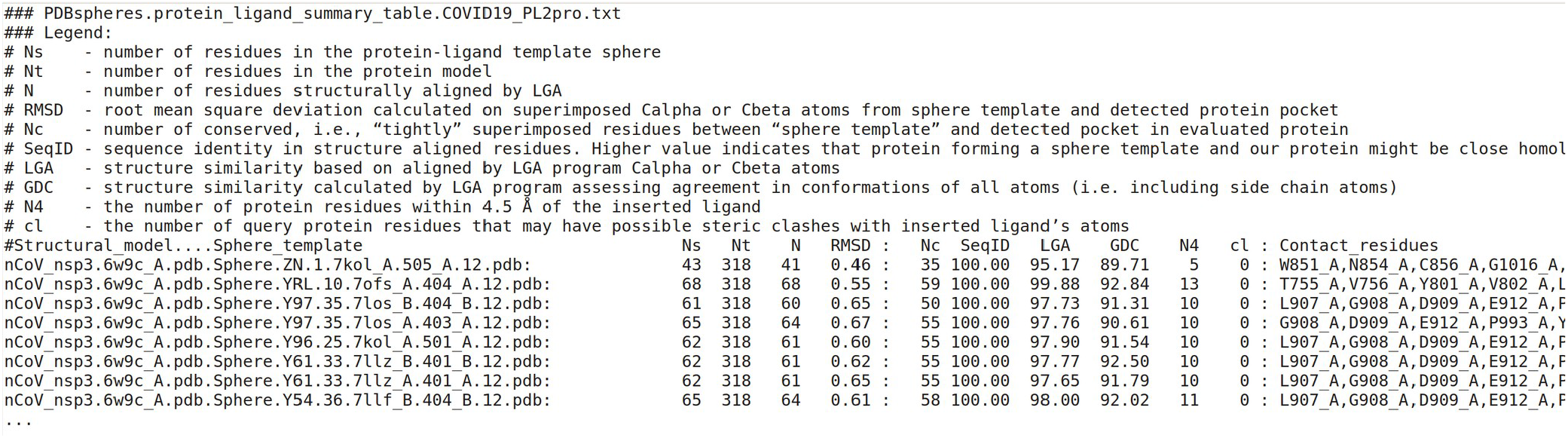
PDBspheres.COVID19_PL2pro.protein_ligand_summary.txt.

**File2:**
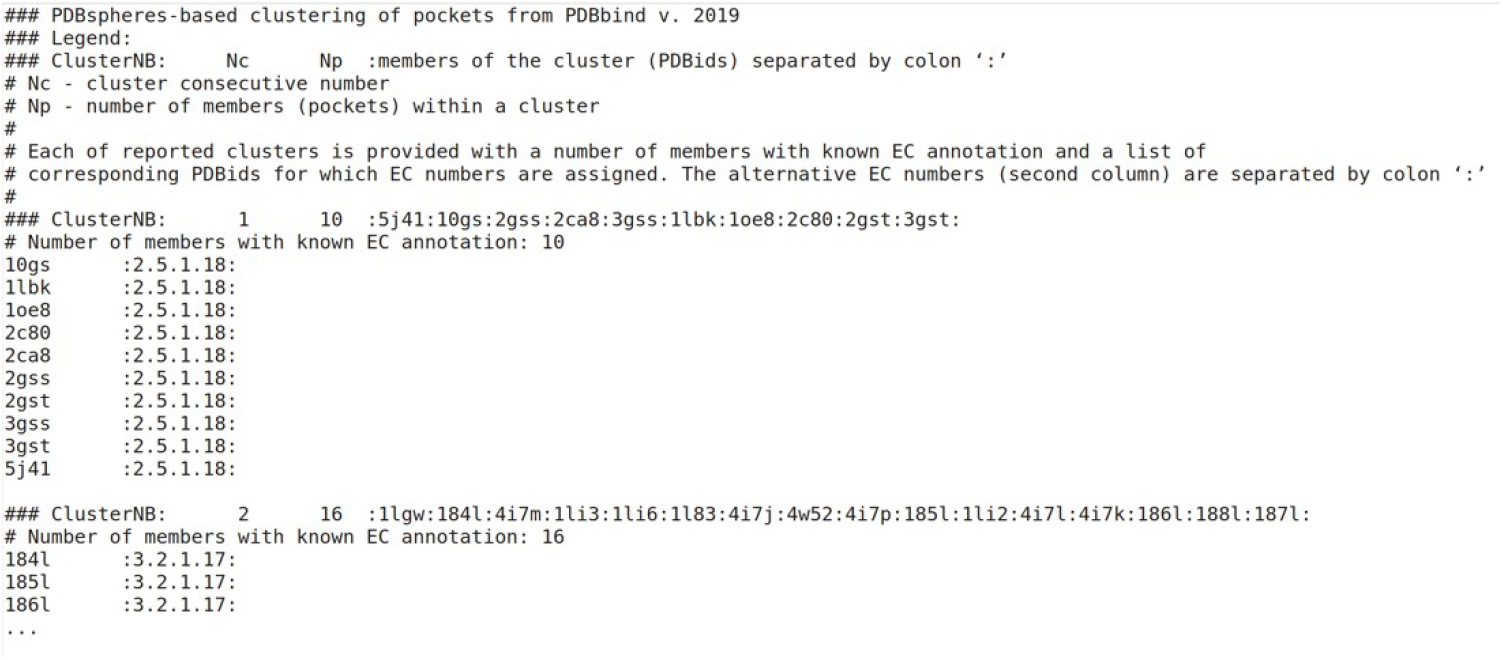
PDBspheres.PDBbind_Clusters.EC_included.txt.

**File3: PDBspheres.PDBbind_Clusters.SCOP_included.txt**

(the same format as File2, but reporting SCOP annotation)

**File4: PDBspheres.PDBbind_Clusters.GO_included.txt**

(the same format as File2, but reporting GO annotation)

**File5:**
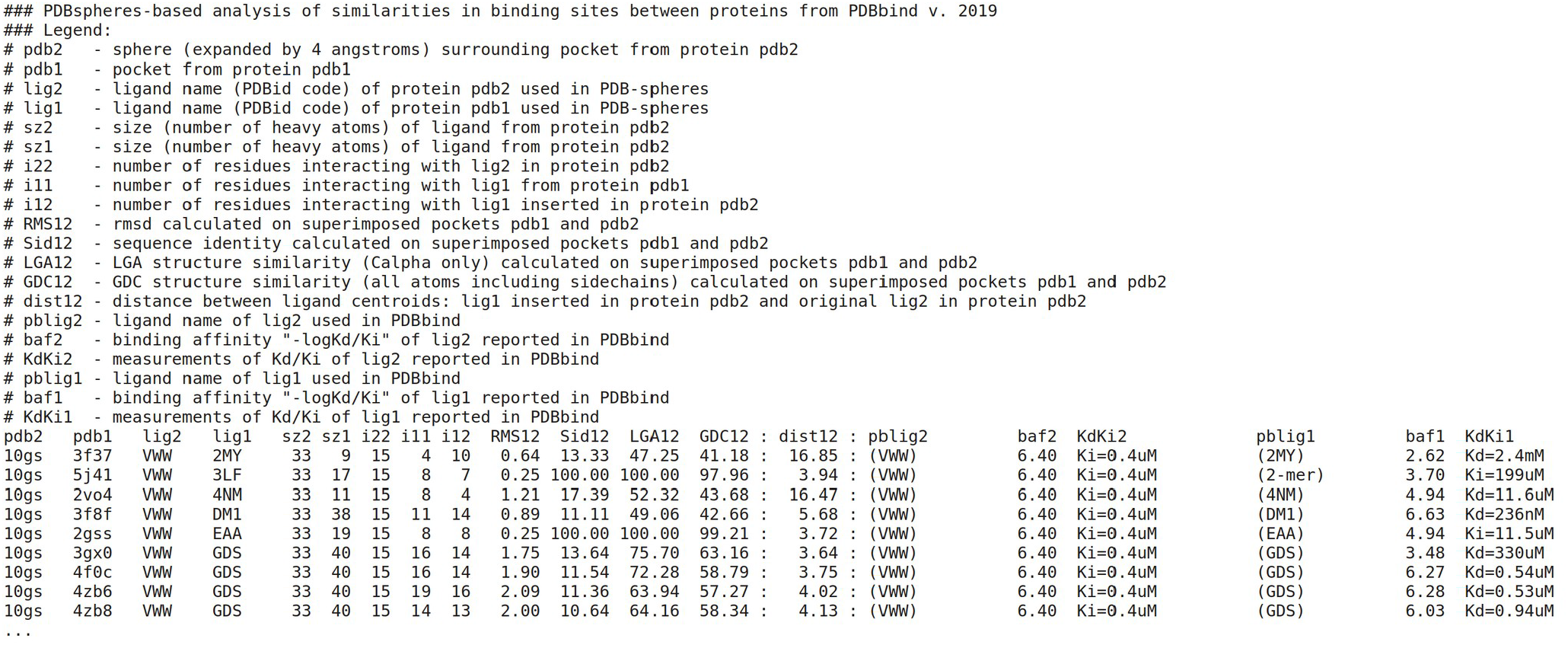
PDBspheres.PDBbind_Binding_sites_similarities.GDC_and_affinities.txt.

**File6: PDBspheres.PDBbind_Clusters.interactive_plot.html (**interactive overview of predicted clusters shown on Figure.7.)

